# Two temperature-dependent membrane fluidity regimes in Gram-positive bacteria

**DOI:** 10.64898/2026.01.22.701129

**Authors:** Aurélien Barbotin, Dimitri Juillot, Paprapach Wongdontree, Rut Carballido-López

## Abstract

It is widely accepted that bacterial cells maintain a constant membrane fluidity in response to temperature changes. This process, known as membrane fluidity homeostasis, occurs through remodeling of plasma membrane composition. We tested this using an assay based on total internal reflection-fluorescence correlation spectroscopy (TIR-FCS) that directly quantifies membrane fluidity as the diffusion speed of a membrane marker in *Bacillus subtilis* and two other Gram-positive bacteria, *Streptococcus pneumoniae* and *Staphylococcus aureus*, across a temperature range of 20°C to 37°C. Instead of the expected constant membrane fluidity, we identified a two-component regime: membrane fluidity is maintained at low temperatures (<26°C), but freely increases at higher temperatures.

**Significance statement:** Temperature changes affect the physical properties of the bacterial plasma membrane. Typically, with a reduction in temperature comes a loss of membrane fluidity. It is known that bacteria like Bacillus subtilis adapt their membrane composition when this happens, and it was thought to maintain membrane fluidity constant. We found that this is not true: fluidity is maintained only in a low (<26°C) temperature regime, while at higher temperatures fluidity is not maintained. We found this to be true for several Gram+ bacteria.

## Introduction

Membrane fluidity changes have been linked to a wide variety of processes in bacteria, for example responses to physical or chemical stresses, protein folding, respiration^1^ and antibiotic resistance^2^. One well-studied aspect of membrane fluidity in bacteria is membrane fluidity homeostasis in response to changes in temperature. In Gram-negative bacteria like *E. coli*, membrane fluidity homeostasis was measured as a gradual increase of membrane FA unsaturation as temperature decreased ^3^. The mechanism of adaptation in Gram-positive bacteria like *B. subtilis*^4^ is more complex: changes of temperature are detected by the two-component system DesK/DesR, which initiates the transcription of the FA desaturase Des upon cold shock^5^. Long-term membrane adaptation involves FA branching and shortening^4^. Membrane FA analysis has long remained the gold standard to evaluate membrane fluidity homeostasis, despite growing evidence of the role of other actors like lipid headgroups ^6^ or proteins ^7^ on the physical properties of the membrane. Another standard approach, which consists in using environment-sensitive probes such as diphenylhexatriene (DPH) to probe local membrane properties, produced conflicting results ^8^. This is not entirely surprising, as environment-sensitive probes report on complex signals ^9^ that correlate with membrane fluidity only under specific biophysical assumptions that are often not verified in biological experiments ^10^.

Recent quantitative measurements revealed that plasma membrane fluidity in *B. subtilis* is approximately two-fold lower at low temperatures (20-23°C) than at 37°C^11,12^. This is in disagreement with the substantial literature that uncovered mechanisms of thermoadaptation in this bacterium. Here, we sought to resolve this apparent contradiction using a technique based on total internal reflection-fluorescence correlation spectroscopy (TIR-FCS) that we recently developed^12^ to directly measure membrane fluidity at a range of temperatures in three different Gram-positive bacteria. We found two distinct operating regimes: at low temperatures (< 26°C), membrane fluidity is maintained constant by the established homeostatic feedback mechanisms; in contrast, at higher temperatures, membrane fluidity increases linearly with temperature.

## Results

We quantified membrane fluidity as the diffusion speed of the membrane marker Nile Red in live wild-type *B. subtilis* cells using TIR-FCS^12^ (Fig. 1A). We first measured membrane fluidity shortly (25-70 mins, after complete cold shock recovery^12^) after cells were transferred from their growth temperature (37°C) to a target temperature between 20°C and 37°C. To our surprise, we observed that membrane fluidity in *B. subtilis* was neither constant nor proportional to temperature. Instead, we observed two operating regimes: membrane fluidity was constant below 26°C, and evolved linearly with temperature above this temperature (Fig 1B). We then verified that removing a gene of the well-studied thermosensing pathway of *B. subtilis* could alter this phenotype. These experiments being performed shortly after the change in temperature, we decided to remove the fatty acid desaturase Des, the fast responder known to immediately desaturate membrane fatty acids after a cold shock^4^. We measured membrane fluidity in a Δ*des* mutant across the same temperature range applied to wild-type cells. We observed that the fluidity was no longer constant below 26°C (Fig. 1C), confirming the role of the Des enzyme in membrane fluidity homeostasis at low temperatures. Using membrane FA extraction and analysis, we measured the relative proportion of the most varying FA between exponentially growing cells at 37°C and 45 mins after a cold shock at 22°C. The only notable difference between both strains was the apparition of unsaturation in WT cells after cold shock explaining the different phenotype of both strains at low temperature. We observed in both strains a similar reduction in overall carbon chain length (Fig. 1D) after cold shock, as previously observed^13^, which was likely the beginning of the long-term membrane fluidity adaptation.

**Figure 1.**
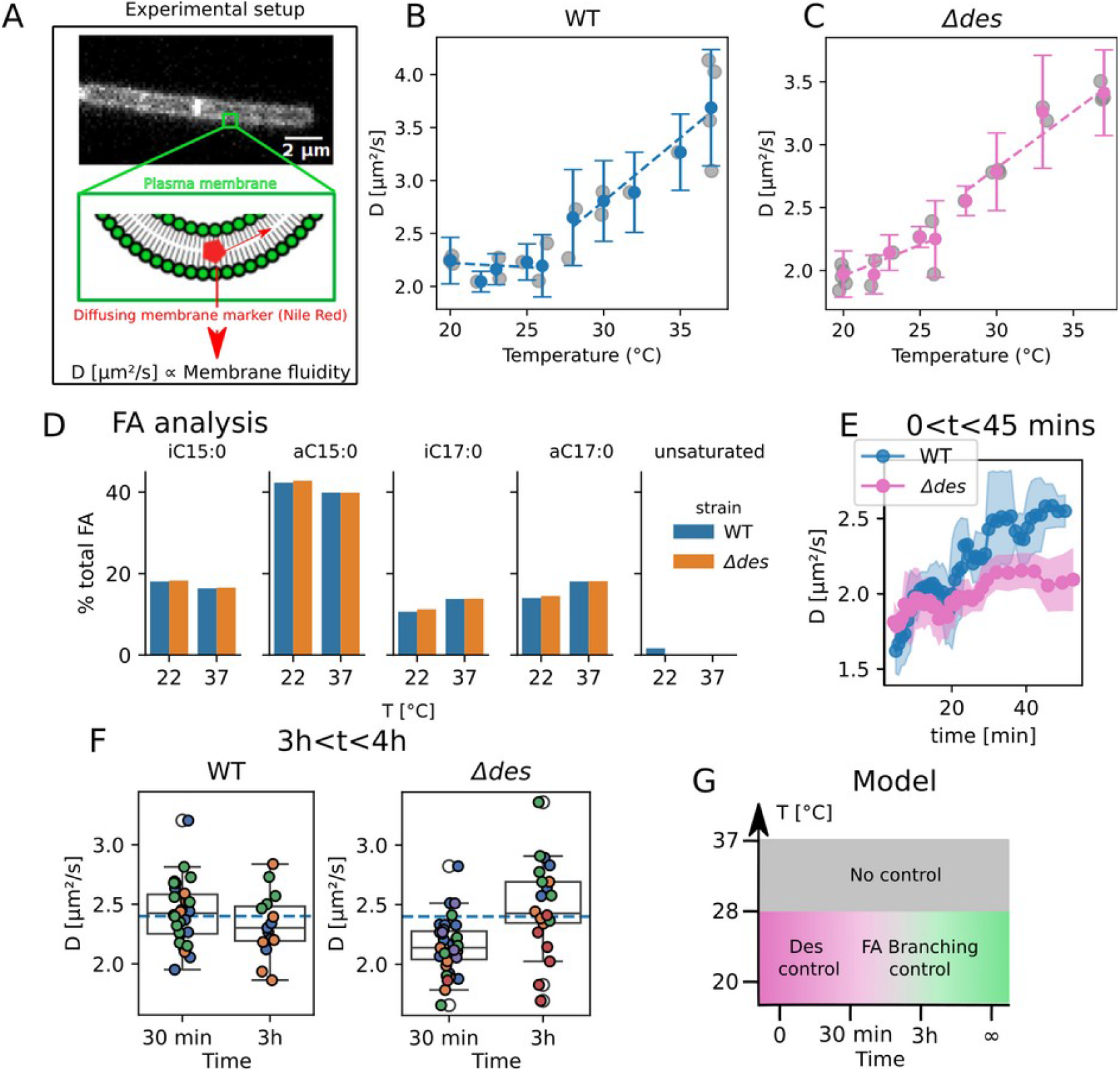
Temperature-dependent membrane fluidity homeostasis in *B. subtilis*. (A) schematic of the methodology. Membrane fluidity is measured as the diffusion coefficient of the membrane marker Nile Red in the plasma membrane. (B-C) Membrane fluidity (mean +/-s.d, gray dots: median diffusion coefficients of individual replicates) in wild-type (WT) (B) and *Δdes B. subtilis* cells (C) measured 25-70 mins after transfer at the target temperature. (D) Relative FA composition of WT and *Δdes* strains at 37°C and 45 mins after cold shock at 22°C. FA types represented are iso (iC) or anteiso (aC) with saturated carbon chains of length 15 or 17, and unsaturated FA (mean of n=2). (E) Time-dependent membrane fluidity recovery in WT (blue) and *Δdes* (pink) cells after a transfer from 37°C to 20°C. (F) steady-state membrane fluidity in WT (left) and *Δdes* (right) ~30 min after cold shock or ~3h after cold shock. (G) Proposed model of temperature-dependent membrane fluidity control in *B. subtilis*.

Measuring the dynamics of cold shock recovery in both wild-type and *Δdes* cells, we observed as expected a much lower recovery of membrane fluidity with time in the Δ*des* mutant than in the WT (Fig. 1D). From this, we concluded that the Δ*des* mutant loses its membrane homeostasis phenotype at low temperatures because of its inability to quickly increase its membrane fluidity. We verified that this inhibition was transient, as predicted by the literature that describes long-term membrane homeostasis to be performed by FA branching and shortening and not unsaturation. For this, we compared membrane fluidity measurements in WT and Δ*des* strains, immediately after cold shock and after approximately 3 hours (~ 1 generation) spent at 20°C. While WT cells exhibited the same membrane fluidity regardless of time spent at 20°C, Δ*des* cells recovered the WT fluidity after 3 hours at 20°C (Fig. 1E). Membrane fluidity of WT cells remained comparatively lower at 20°C than at 37°C even for much longer times spent at 20°C: the diffusion coefficient of Nile Red remained below 2.5 µm^2^/s even in cells that grew for 24h at 20°C (data not shown). These observations led us to propose a new model of membrane fluidity homeostasis, to complete the current model and reconcile conflicting observations (Fig. 1F). Our observations do not contradict that the two mechanisms uncovered so far of FA unsaturation and branching are indeed activated at different timescales after a cold shock to maintain membrane fluidity. However, our data revealed that above a critical temperature, which we estimate around 26°C (Fig. 1B), membrane fluidity is not controlled.

To verify whether this model is specific to *B. subtilis* or extends to other Gram-positive bacteria, we performed TIR-FCS temperature scans in two Gram-positive mesophilic pathogens, *Staphylococcus aureus* and *Streptococcus pneumoniae* (Fig. 2). These data clearly show that both bacterial species exhibit a membrane fluidity profile similar to that observed in *B. subtilis*: membrane fluidity remains constant below a certain critical temperature, and increases linearly with temperatures above this threshold. The exact value of this critical temperature seems to be around 26°C in both *S. pneumoniae* and S. aureus as in *B. subtilis*, yet the noisier TIR-FCS measurements in these two species precludes drawing a definitive conclusion. Notably, the diffusion coefficients across all three bacteria are remarkably similar (ranging from 2-2.5 µm^2^/s at lower temperatures to ~3.5 µm^2^/s at higher temperatures). This could hint at a conserved membrane fluidity control mechanism across all three bacteria, in accordance with recent findings that identified thermosensor proteins similar to those of B. subtilis in *S. aureus* and *Bacillus anthracis* ^14^.

**Figure 2.**
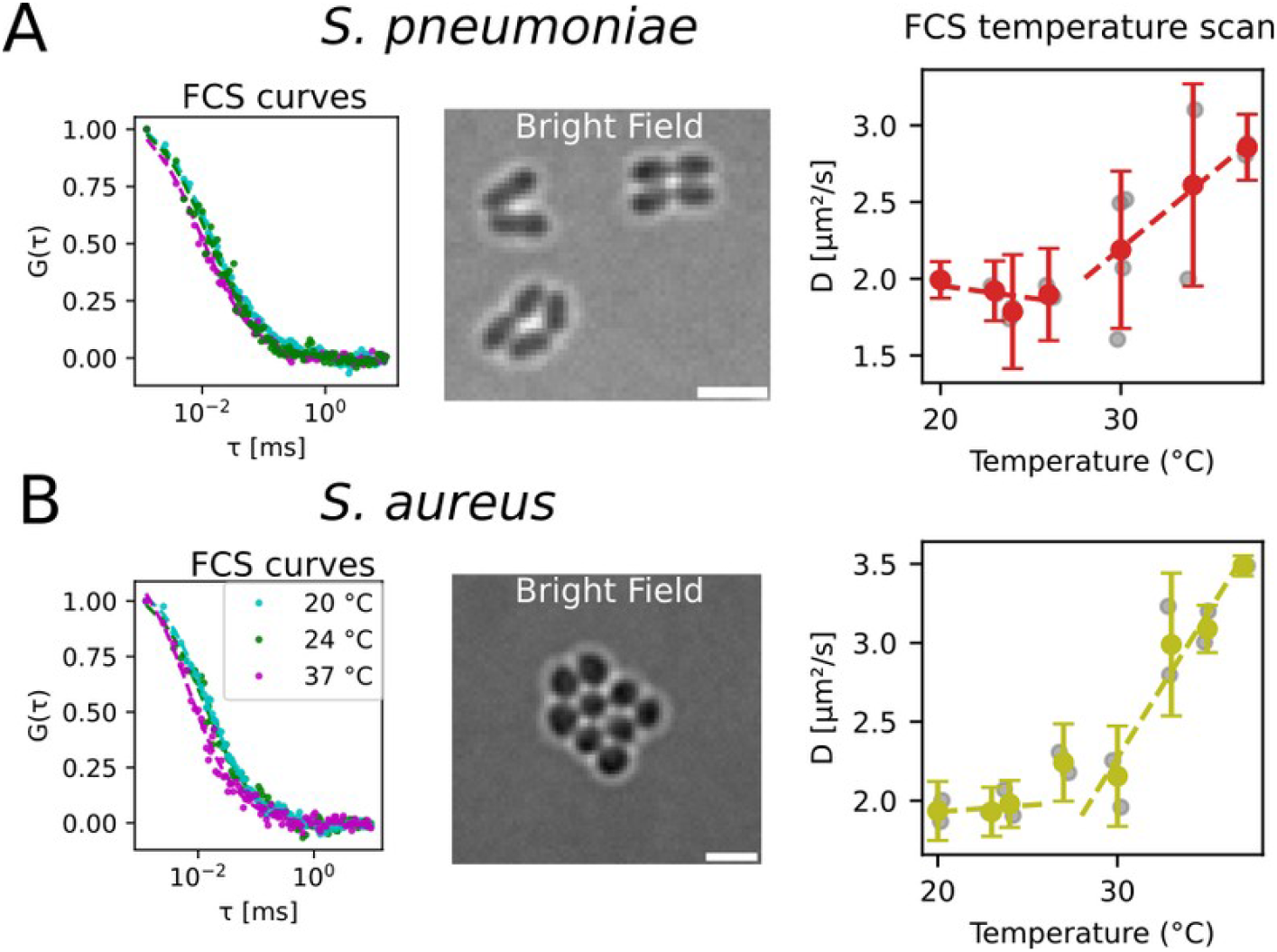
FCS temperature scans in *S. pneumoniae* (A) and *S. aureus* (B). Left, Representative FCS curves at different temperatures. Middle, bright-field images (scale bar: 2 µm). Right, temperature-dependent membrane fluidity.

## Discussion

Through quantitative measurements of membrane fluidity, we uncovered the existence of two distinct operating regimes of membrane fluidity in several Gram-positive bacteria, instead of one as previously thought. This discovery does not contradict previous data, in particular measured changes in membrane composition as a response to cold, but rather gives it a new perspective. The temperature threshold between the two regimes, which we estimate to be around 26°C in *B. subtilis* based on our data, is in line with the previously reported activation temperature of the thermosensor desK ^15^.

We believe that our finding raises many interesting questions. What is the point of these two operating regimes? Based on membrane FA analysis^3^, it is likely that Gram-negative bacteria do not exhibit this behaviour in this range of temperatures: why and what does this imply for the functions regulated by membrane properties? We observed the same regimes in three Gram-positive mesophilic bacteria, but what about psychrophilic bacteria? In *B. subtilis*, the thermosensor desK is thought to measure membrane thickness and not fluidity directly ^4^. We can reasonably expect that membrane thickness will follow the same two-component behaviour and it is likely to be the case for many other physical parameters of the membrane. Finally, this work highlights that caution is required when quantifying motion of membrane-associated objects at physiological temperatures, as minor changes in temperatures can lead to substantial measurement bias. When the highest precision is required, higher stability can be attained by working in the membrane-fluidity controlled regime.

## Materials and Methods

Diffusion speed of membrane marker Nile Red was measured with TIR-FCS^12^. Each acquisition of 50 000 frames was about one minute long. Raw diffusion coefficients were divided by a factor specific to the shape of each bacterium (*B. subtilis*: 1.54, *S. aureus* 2.38 and *S. pneumoniae* 2.19) estimated from simulations. Detailed methods can be found in the appendix.

## Acknowledgements

We thank Drs Charlène Cornilleau and Armand Lablaine and the members of the ProCeD laboratory for their support and helpful discussions. We also thank Dr Daniela Albanesi for useful discussion. We thank Drs Alexandra Gruss and Florence DUbois-Brissonnet for their help with FA extraction and analysis. This project was funded by a Consolidator grant from the European Research Council (ERC) under the Horizon 2020 research and innovation program (ERC-2017-CoG-772178 to R.C.-L.) and under the Marie Sklodowska-Curie grant agreement no. 101030628.

