## extended methods for "Two temperature-dependent membrane fluidity regimes in Gram-positive bacteria"

### TIR-FCS

TIR-FCS acquisitions consisted of 50000 frames, with a frame interval time to 1.26 ms, maximum achievable speed on our setup. Pixel size was set to 160 nm and the pixels of resulting TIR-FCS stacks were binned 3 by 3 to reduce noise, leading to an effective pixel size of 480 nm. FCS curves were fitted with a 2D diffusion model model, accounting for the bias induced on diffusion coefficient measurement by the high cellular curvature. We estimated the bias using simulations, as described in ref. [12] . In the case of S. pneumoniae, exhibiting two different curvatures alongside the long and short axis, we estimated the bias as

$bias=\sqrt{\square}$

We considered an average cell diameter in WT *S. pneumoniae* of 700 nm and an average cell length of 1.5 µm, an average cell width and length of 1 and 3 µm in B. subtilis and an average cell diameter of 1 µm in S. aureus. Simulating these morphologies yielded the corresponding correction factors:

| B. subtilis | S. aureus | S. pneumoniae |
|---|---|---|
| 1.54 | 2.38 | 2.19 |

These factors are slightly higher than the ones we previously used in ref. [12] due to the larger effective pixel size we used in this study.

TIR-FCS imaging was performed on cells immobilised on agarose pads. Cells were left to accommodate on the pad for at least 20 mins prior to imaging.

Excitation laser at 561 nm was set to a power of approximately 70 nW/µm² in *B. Subtilis* and *S. aureus*, and 140 nW/µm² in *S. pneumoniae*, corresponding to respectively 5 and 10 % of the maximum power available to this laser. Acquisition speed was set to 1.26 ms per frame, the maximum achievable on our system.

#### *Bacillus subtilis* cultures

Cells were grown overnight at 30°C in LB. They were diluted in the morning in LB to an OD of about $6.10^{-3}$. Cells were grown for approximately 2h30 to an early exponential phase corresponding to an OD of 0.1-0.3.

#### *Staphylococcus aureus* cultures

Cells were grown in LB, overnight at 37°C, then diluted in the morning in LB to an OD of about $1.10^{-1}$. Cells were grown for approximately 2h to an early exponential phase corresponding to an OD of 1.

#### *Streptococcus pneumoniae* cultures

After thawing stock cultures, aliquots were inoculated at $OD_{550}$ of $6.10^{-3}$ in C+Y medium (reference : 10.1007/978-1-4939-9199-0_6 ) and grown at 37°C to an $OD_{550}$ of 0.3. These precultures were then inoculated (1/50) in C+Y medium and incubated at 37°C to an $OD_{550}$ of 0.15.

### Sample preparation for microscopy

Cells were labeled with 0.2% of a solution of Nile Red at a stock concentration of 20 µg/mL in DMSO for 10-20 mins under agitation. Cells were then immobilised on an agarose pad made of LB and 1.2% agarose. A freshly plasma-cleaned coverslip was added. For FCS temperature scans, cells were immobilised on pad and immediately transferred to the target temperature for at least 20 mins before the start of the experiment, except in *B. subtilis* at 20°C where cells were first left to accommodate on pad at 37°C for 20 mins then transferred at 20°C for both cold shock recovery measurements and later steady-state diffusion measurements at 20°C. No cell spent more than 70 mins on pad.

### Data processing and visualisation

Each acquisition of 50000 frames was performed as follows. Pixels containing cells were semi-manually selected. FCS curves in these pixels were fitted and selected for analysis. If less than a given threshold of FCS curves was considered acceptable by our quality metric (ref. [12]), we considered the acquisition to be artefactual and was discarded. This threshold varied with bacterium, as data quality was uneven between bacterial species. This threshold was set to 70 % of valid curves for B. subtilis and to 40% for the others. The median diffusion measurement from each acquisition was considered a data point. For a replicate, typically 6 acquisitions were performed corresponding to 6 data points. The average of these datapoints represent the average of a replicate, represented by gray points in Figs. 1B-C and Fig. 2. Colored points and error bars correspond respectively to mean and standard deviation of all data points at a given temperature. Trend lines were drawn by fitting two lines to the data, one for the points between 20 and 26°C and one between 28 and 37°C.

### Fatty acids extraction

*B. subtilis* cultures were harvested at an $OD_{600}$ of ~0.3. Bacterial cultures were frozen in liquid nitrogen directly. Cells were centrifuged and washed once with 0.9% NaCl. Cell pellets were treated with 500 µL of 1 N sodium methoxide in methanol (reference: 10.1016/j.celrep.2019.11.071). Subsequently, 200 µL of heptane containing methyl-10-undecenoate (Sigma-Aldrich) as the internal standard was added, vortexed for 1 min, and rapidly centrifuged at high speed. Fatty acid methyl esters were collected in the heptane phase. Analyses were performed on an ISQ 7000 Single Quadrupole Gas Chromatograph (Thermo Scientific). Data were acquired and processed using Chromeleon 7.3 (Thermo Scientific). *B. subtilis* fatty acid peaks were detected and identified by MS library search. The average of two biological replicates is shown for each condition, and only the FA exhibiting the largest relative changes between conditions are shown for the sake of concision.

### Strains

We used *B. subtilis* WT strain 168 (laboratory stock) and Δ*des*::*km* strain RCL0844 (https://doi.org/10.1128/mSystems.01017-2152,139).

We used *S. aureus* attenuated WT strain RN-4220 ( https://doi.org/10.1128/mbio.03193-20103,254)

We used *S. pneumoniae* strain R1501 derived from the WT R800 (https://doi.org/10.1111/j.1365-2958.2003.03892.x )
